# The abundance change of age-regulated secreted proteins affects lifespan of *C. elegans*

**DOI:** 10.1101/2024.08.16.608229

**Authors:** Prasun Kumar Bhunia, Vishwajeet Raj, Prasad Kasturi

## Abstract

Proteome integrity is vital for survival and failure to maintain it results in uncontrolled protein abundances, misfolding and aggregation which cause proteotoxicity. In multicellular organisms, proteotoxic stress is communicated among tissues to maintain proteome integrity for organismal stress resistance and survival. However, nature of these signalling molecules and their regulation in extracellular space is largely unknown. Secreted proteins are induced in response to various stresses and aging, indicating their roles in the inter tissue communication. To study fates of age-regulated proteins with potential localization to extracellular, we analysed publicly available age-related proteome data of *C. elegans*. We found that abundance of proteins with signal peptides (SP) increases with age and result in their aggregation. Intriguingly, these changes are differentially regulated in the lifespan mutants. A subset of these SP proteins is also found in the cargo of extracellular vesicles. Many of these proteins are novel and functionally uncharacterized. Reducing levels of a few extracellular proteins result in increasing lifespan. This suggest that uncontrolled levels of extracellular proteins might disturb proteostasis and limit the lifespan. Overall, our findings suggest that the age induced secreted proteins might be the potential candidates to be considered as biomarkers or for mitigating age-related pathological conditions.

## 1. Introduction

With advancing age, cells encounter challenges in properly folding, assembling, and degrading proteins, leading to imbalance in protein homeostasis or proteostasis (Hipp et al., 2019). This imbalance often results in the accumulation of misfolded or aggregated proteins, which can have detrimental effects on cellular function and contribute to the pathogenesis of various age-related diseases, including neurodegenerative disorders like Alzheimer’s, Parkinson’s, and Huntington’s diseases (Hipp et al., 2014). Accumulation of extracellular protein aggregation of proteins such as beta-amyloid, tau, alpha-synuclein is also a hallmark feature of these disorders (Jankovska et al., 2020; Rahman and Lendel, 2021). Additionally, evidence suggests that extracellular aggregates can propagate between cells, spreading pathology in a prion-like manner Brundin et al., 2010). Inside the cells the protein quality control systems include molecular chaperones that assist in protein folding and preventing misfolding and the ubiquitin-proteasome/autophagy systems degrade damaged or excess proteins (Klaips et al., 2018; Sinnige et al., 2020). Similarly, protein quality control mechanisms in the extracellular space are vital for maintaining extracellular proteostasis and facilitating inter-tissue communication (Takeuchi, 2018; Wilson et al., 2023). However, less is known how the proteostasis is maintained in the extracellular space.

In multicellular organisms, different cells and tissues communicate and coordinate in maintaining the proteome integrity under various conditions (Sala et al., 2017). Secreted proteins serve as vital mediators of inter-tissue communication and maintaining homeostasis throughout the organism. These proteins, synthesized and released by one cell type, often act upon distant targets to convey signals that regulate cellular functions and metabolism. Once proteins are secreted from cells into the extracellular environment, they are subjected to various challenges that may compromise their structure and function (Mesgarzadeh et al., 2022). The precise control of secreted protein abundances, activity, and localization is essential for the proper inter-tissue communication. Emerging evidences suggest that protein quality control in the extracellular space include a few proteins that function as chaperones and proteases (Satapathy and Wilson 2022; Takeuchi et al., 2015). Furthermore, exosomes or extracellular vesicles also play a role in protein quality control by transporting proteins to specific destinations or facilitating their degradation (Takeuchi et al., 2015; Wang and Barr, 2018). Together, they maintain overall quality of the extracellular proteome which is crucial for the integrity and functionality of secreted proteins, ultimately contributing to the organismal homeostasis. Dysregulation of secreted proteins and their signalling pathways can lead to pathological states, including metabolic disorders and age-related maladies (Kuo et al., 2021; Takeuchi, 2021). Thus, understanding the mechanisms underlying inter-tissue communication mediated by secreted proteins is critical for elucidating physiological processes implicated in aging such as organism wide proteostasis.

Cell-nonautonomous regulation of health, longevity and organismal proteostasis has been reported in model organisms (O’Brien and van Oosten-Hawle, 2016; Miller et al., 2020; Morimoto, 2020). It has been observed that the abundance of putative extracellular signalling proteins (secreted proteins including neuro peptides) increases during stress and aging (Liang et al., 2014; Walther et al., 2015; Narayan et al., 2016). Many of these secreted proteins are functionally uncharacterized (Suh and Hutter, 2012), raising the question, whether they play roles in extracellular proteostasis and inter-tissue stress signalling. Identification, characterization and modulation of these signalling molecules and the pathways thereof will help in understanding the molecular mechanisms of the inter-tissue communication in maintaining organism wide proteostasis that improve healthy lifespan. Ultimately it may lead to design of future therapeutic interventions for age- and other stress-associated maladies.

In *C. elegans*, the extracellular proteins directly secreted to body fluid which bathes the internal organs, making it an attractive system to study and identify extracellular proteins. Here we report identification of age-related changes in the proteins that contain signal peptides and potential to be localized to extracellular. We systematically looked at their abundance levels and aggregation during aging in wildtype and lifespan mutant worms. Some of these proteins are also associated with extracellular vesicles. We found that abundance of some of these proteins increases continuously during aging and are differentially regulated in long and short lifespan mutant worms. Reducing levels of a few of these proteins results in lifespan increase, suggesting high abundance of these proteins contribute to proteostasis imbalance and limit the lifespan of the worms.

## 2. Materials and methods

### 2.1. Data collection and analysis

Aging proteome of *C. elegans* was downloaded from the published datasets by Narayan et al and Walther et al (Walther et al., 2015; Narayan et al., 2016) and analysed using Perseus_2.0.6.0 program (Tyanova et al., 2016). The Narayan dataset contains 6295 and the Walther dataset contains 4078 proteins (based on at least three valid values). There are 3061 proteins that are common to both datasets. We screened out the proteins containing signal peptides using SignalP-6.0 (Teufel et al., 2022) and identified their subcellular localization by CELLO v. 2.5 web server (Yu et al., 2006; Yu et al., 2014). We found 738 and 607 SP proteins in Narayan and Walther datasets respectively and 429 SP proteins are common in both datasets. The proteins were sorted according to their fold change (as *log2* ratios) and gene ontology (GO) analysis was performed using Shiny GO 0.80 webserver (Ge et al., 2020). Protein-protein interaction (PPI) network analysis was performed using Metascape (Zhou et al., 2019). Extracellular vesicle (EV) proteome data was from downloaded from the published datasets by Russell et al and Nikonorova et al (Russell et al., 2020; Nikonorova et al., 2022). Protein abundance changes in WT and lifespan mutants (*daf-2* and *daf-16*) were calculated as fold changes at indicated time points (days) and comparisons were made relative to day1 or as mentioned in the results section.

### 2.2. C. elegans maintenance

The N2 wild type strain from the Caenorhabditis Genetics Center (CGC) was used for this study. Worms were grown on NGM plates with OP50 bacteria and maintained at 20°C as described (Stiernagle et al., 2006).

### 2.3. Lifespan analysis

The feeding RNAi method was used to knock down expression of the selected candidate genes. Empty vector (L4440) was used as control. HT115 bacteria carrying the RNAi clones were grown on NGM plates containing IPTG (1mM) and carbenicillin (25mg/ml). All RNAi clones were confirmed by sequencing. The worms were synchronized by egg laying method. The wildtype adult worms were added to the RNAi plates and L4 worms from the progeny were transferred to new RNAi plate. Next day, after laying eggs (for 4hrs), the worms were removed to get synchronized population. 100 to 120 worms were used per lifespan assay. First day of the adulthood was considered as day1. Worms were transferred to new plates every alternate day until they stop laying eggs to eliminate progeny. Worms scored dead if they do not respond to gentle poking. Accidental deaths or bag of worms were censored.

### 2.4. Statistical analysis

Survival analysis was performed using OASIS 2 online application (Han et al., 2016). Origin and GraphPad software were used for graphing and statistical analysis. Statistical tests and p-values are mentioned in the figure legends.

## 3. Results

### 3.1. Identification of age-related changes in the potential secreted proteins (SPs)

Secreted proteins are crucial for cell communication, provide structural support, and mediate responses to various stresses. Some of them might act as protein quality control components in the extracellular region (Gallotta et al., 2020). It has been observed that many secreted proteins are induced in response to heat stress, pathogen infections and during aging (Liang et al., 2014; Walther et al., 2015; Narayan et al., 2016; Suh and Hutter, 2012). Why secreted proteins are induced during aging and fate of these proteins is largely unknown. To identity the secreted proteins that change their abundance during aging, we analysed age-related proteome data from *C. elegans* model organism. The two data-sets we analysed were published by Walther et al (Walther et al., 2015) and Narayan et al (Narayan et al., 2016) and are publicly available. First, we sorted out the proteins containing signal peptide (SP) using SignalP 6.0. We identified 738 SP containing proteins from Narayan data and 607 SP containing proteins from Walther data. In total, 916 SP proteins were identified, in which 429 proteins are common in the both datasets (Fig. 1A, Table S1A). Next, we identified subcellular localizations for these 916 SP proteins using CELLO v. 2.5. This program uses four criteria for predicating localization, such as the amino acid composition, the di-peptide composition, the partitioned amino acid composition and the sequence composition based on the physico-chemical properties of amino acids. The results are also presented as neighbor, combined and most-likely-location category. The outcome may vary for each criterion (Table S2A). Under most-likely-location category, we found 403 proteins are predicted to be localized to extracellular. However, there are proteins which are not predicted as extracellular in this category, are actually extracellular (as confirmed on wormbase (Harris et al., 2020)). For example, the five *ule* genes codes for secreted proteins (Zimmerman et al., 201536), but only ULE-1 (O01780) and ULE-4 (O62053) were predicted as extracellular in the most-likely-location category. However, other ULE proteins (Q067X2, Q18947 and Q9XWT3) were predicted as extracellular in at least one of the other categories. Other examples are *endu-1* (Q21109) and *tag-196* (O16454) which are extracellular (34), but not predicted as by CELLO v. 2.5. So, if any protein is predicted as extracellular in at least one of the four main categories, we considered them as extracellular. With this criterion we obtained 601 proteins (Table S2B). Then we checked the other remaining 315 proteins manually on the wormbase and found that many of them are also extracellular. For example, the five identified VIT (vitellogenin) proteins are secreted/extracellular but were not predicated as extracellular by CELLO v. 2.5. To avoid any discrepancy, we continued to use the 916 proteins for further analysis and called them as SP proteins.

**Fig. 1.**
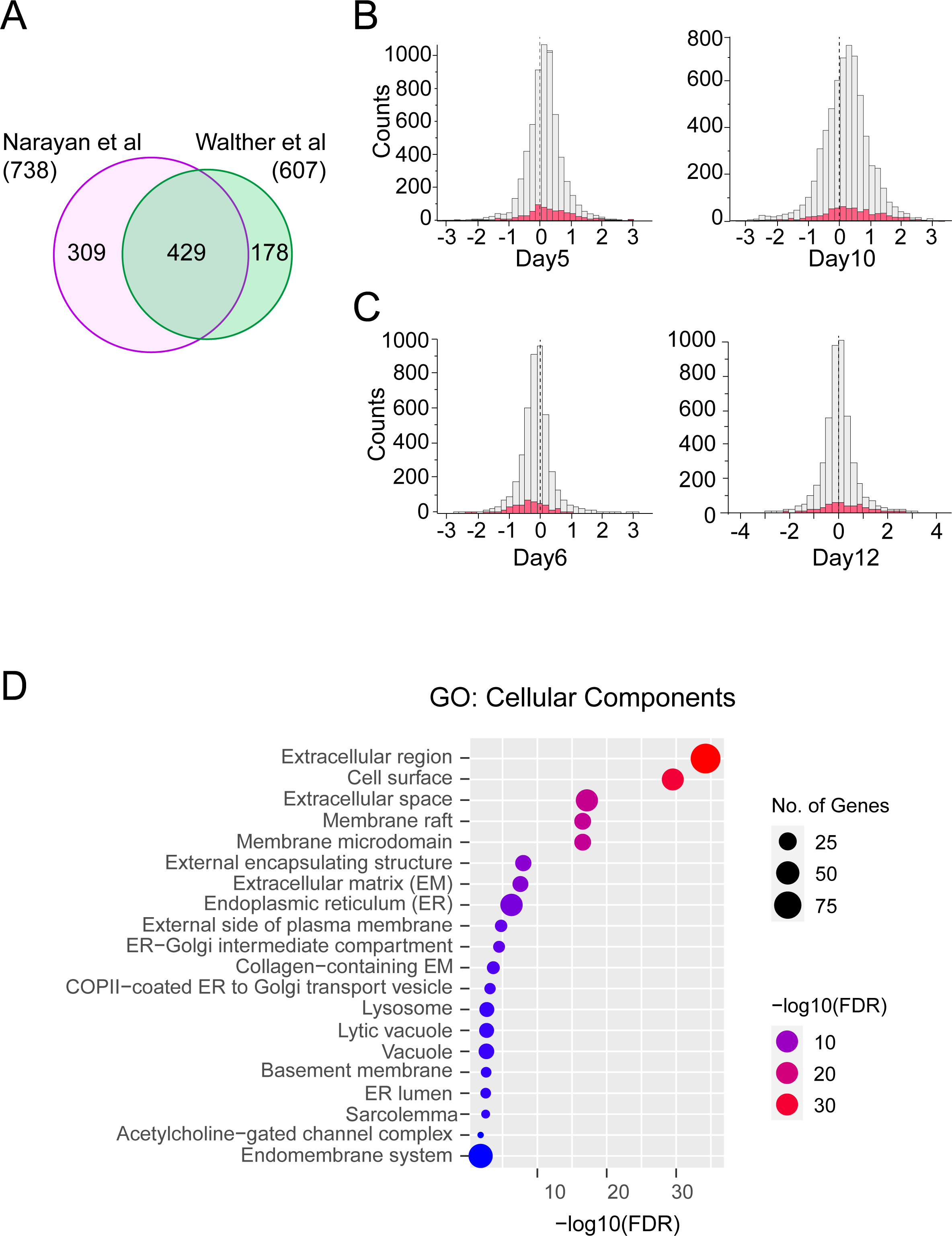
Identification of age-related changes in SP (signal peptide containing) proteins. **A** Venn diagram showing the overlap between SP proteins identified in Narayan and Walther datasets. There are 738 and 607 SP proteins in Narayan and Walther datasets respectively in which 429 SP proteins are common. **B-C** Histograms representing the distribution of the SP proteins in the total proteome of Narayan dataset (day 5 and day10 compared to day1) (B) and Walther dataset (day6 and day12 compared to day1) (C). The x-axis represents log2 ratios at the indicated days for the Narayan and Walther datasets respectively. The light grey color denotes the total number of proteins whereas the red color denotes the SP proteins. **D** Enriched gene ontology (GO) cluster of cellular components (CC) of all the 916 identified SP proteins in this analysis.

To found how these SP proteins change with age, first we looked at their abundance distribution at different times during aging. The abundance levels shifted to higher side with age (Fig. 1B-C). In aged worms (day10 from Narayan data and day12 from Walther data) the abundance increase is more prominent than the decrease with age (Suppl Fig. 1A-B). At day12, in Walther data, 179 and 82 proteins (out of 503) increased and decreased respectively in their abundance more than 2-fold. In Narayan data (at day10), 201 and 90 proteins (out of 738) increased or decreased respectively in their abundance more than 2-fold. Next, we performed GO term analysis for cellular component (CC) and found enrichment of extracellular or components of secreted pathways (Fig. 1D). We also checked only for the up or down regulated proteins and found similar enrichment (Suppl Fig. 1C). Other subcellular compartments are not enriched supporting that the 916 SP proteins might enter ER secretory pathway to reach extracellular region and ER derived organelles or membrane. To further support our findings, we checked published literature and found that some of these SP proteins are indeed secreted/extracellular (Zimmerman et al., 2015; Gallotta et al., 2020).

### 3.2. PPI and network analysis of the SP proteins

To explore more about these SP proteins, we performed protein-protein interactions (PPI) and network analysis using Metascape. First, we analysed all the 916 SP proteins to find how they form network of clusters. We found many protein classes in these networks (Fig. 2A and Suppl Fig. 2A), where some protein families are nematode specific. The number of proteins in each class and the number of identified proteins is listed in the table 1. Many of these SP proteins appear to be uncharacterized, however they might involve in important cellular functions. The PPI analysis identified multiple nodes consisting of such as metabolic, lipid binding, ion transport and protein folding.

**Fig. 2.**
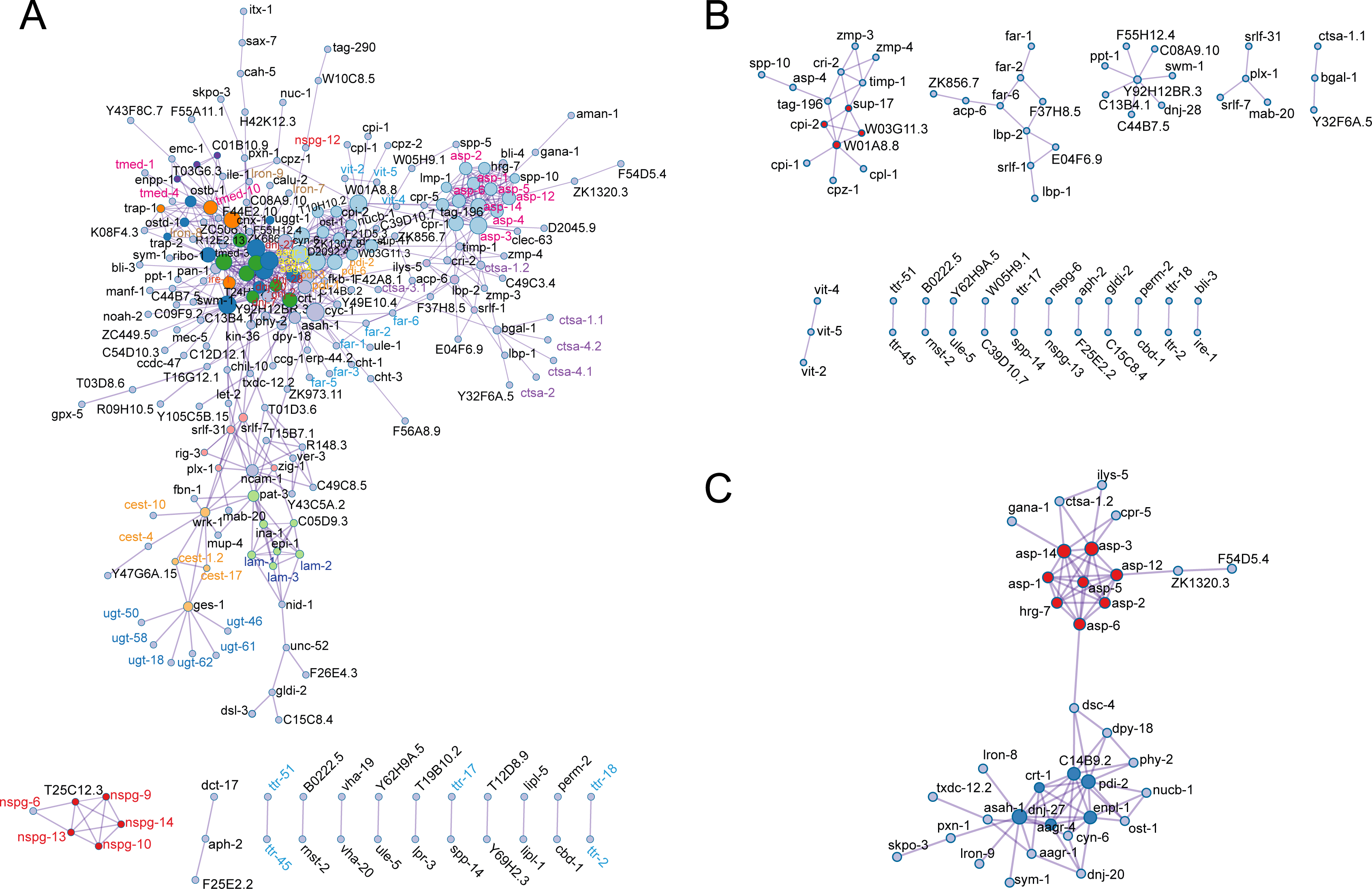
PPI and network analysis of SP proteins. **A** Protein-Protein Interaction (PPI) Network showing the possible interactions and clusters of all the identified SP proteins. **B** PPI for the SP proteins upregulated by 1.5-fold. **C** PPI for the SP proteins downregulated by 1.5-fold.

**Table-1.**
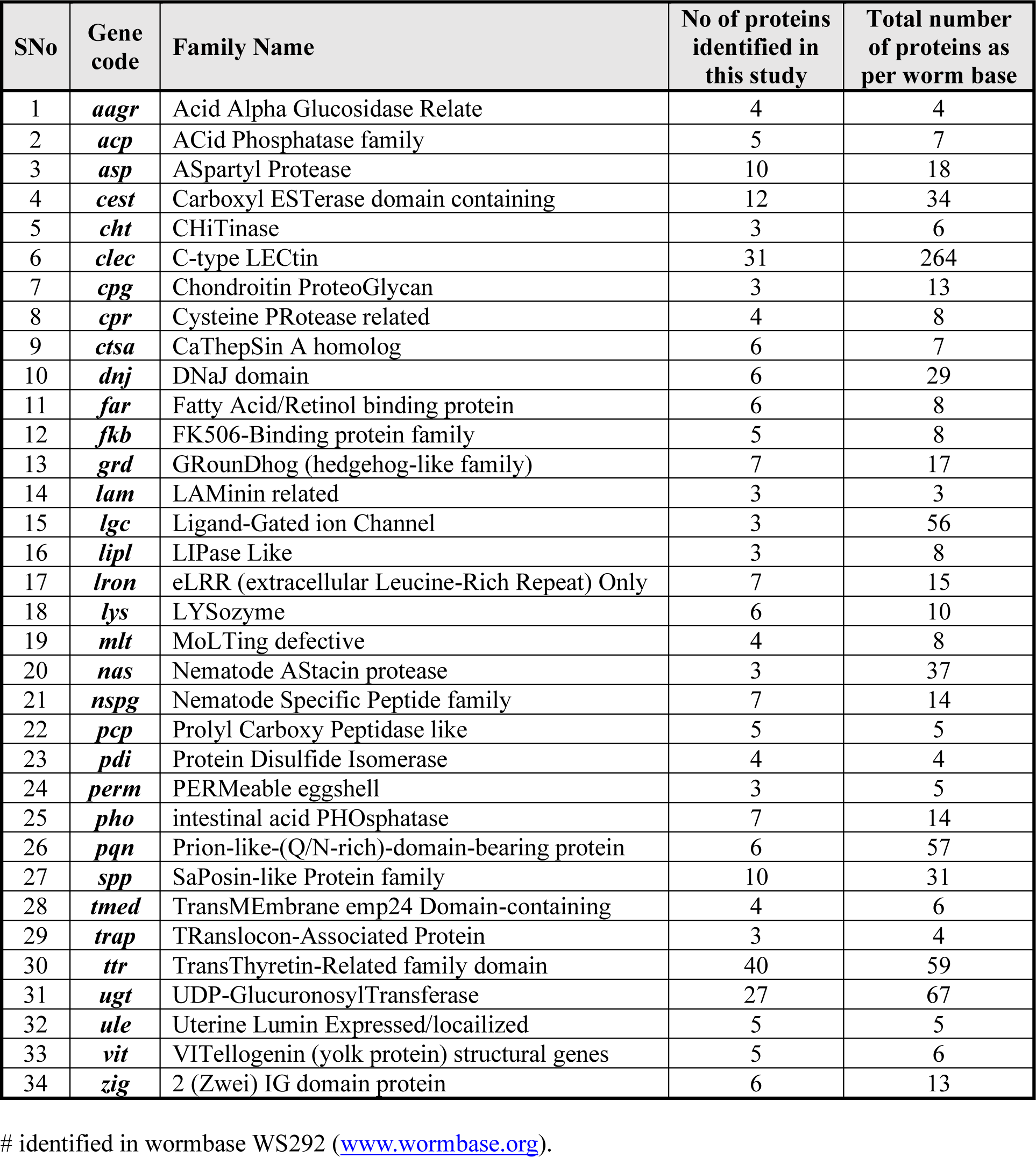
Gene classes identified for SP proteins^#^.

Then, to check how these PPI networks change with age, the SP proteins with abundance increase or decrease by 1.5-fold (on day12) were analysed. Proteins that upregulated with age are also formed similar clusters, however fatty acid binding and endopeptidase inhibitors are among the significantly enriched terms (Fig. 2B and Suppl Fig. 2B). In contrast, proteins that downregulated with age were formed less clusters and enriched endopeptidase activity and unfolded protein binding (Fig. 2C and Suppl Fig. 2C). These results suggest diverse functions for the SP proteins that might alter with age and result in adverse effects.

### 3.3. Abundance of the SP proteins during aging

Since secreted proteins involve in various cellular process including signalling, their abundance change during aging might have consequences on overall health of the organism. To find how the SP proteins abundance change during aging, we checked their fold changes relative to day1 of wildtype worms. We found that abundance increase is more prominent than the decrease which is consistent with both the datasets (Fig. 3A-B, Suppl Fig. 3A-B and Table S3). Proteins that increase their abundance more than 4-fold and 8-fold continue to follow the trend during aging. In Walther data, to the proteins that increase more than 8-fold, the SP proteins contribute to 60% increase (33 of 55) at day6 and 70% increase (73 of 104) at day12 (Fig. 3A). In contrast, proteins that decrease 8-fold, only 38% (8 of 21) come from the SP proteins at day6 and 42% (14 of 33) by day12 (Fig. 3B). To the proteins that increase more than 4-fold, the SP proteins contribute to ∼50% increase (63 of 127) at day6 and 53% increase (117 of 220) at day12 (Supple Fig. 3A). However, in the 4-fold decrease, the SP proteins contribute 49% (24 of 49) at day6 and ∼39% (32 of 81) by day12, which further decreased to 31% by day17 (Suppl Fig. 3B). Similar trend was observed with the Narayan data, however with a smaller number of proteins (Supple Fig 3C-D).

**Fig. 3.**
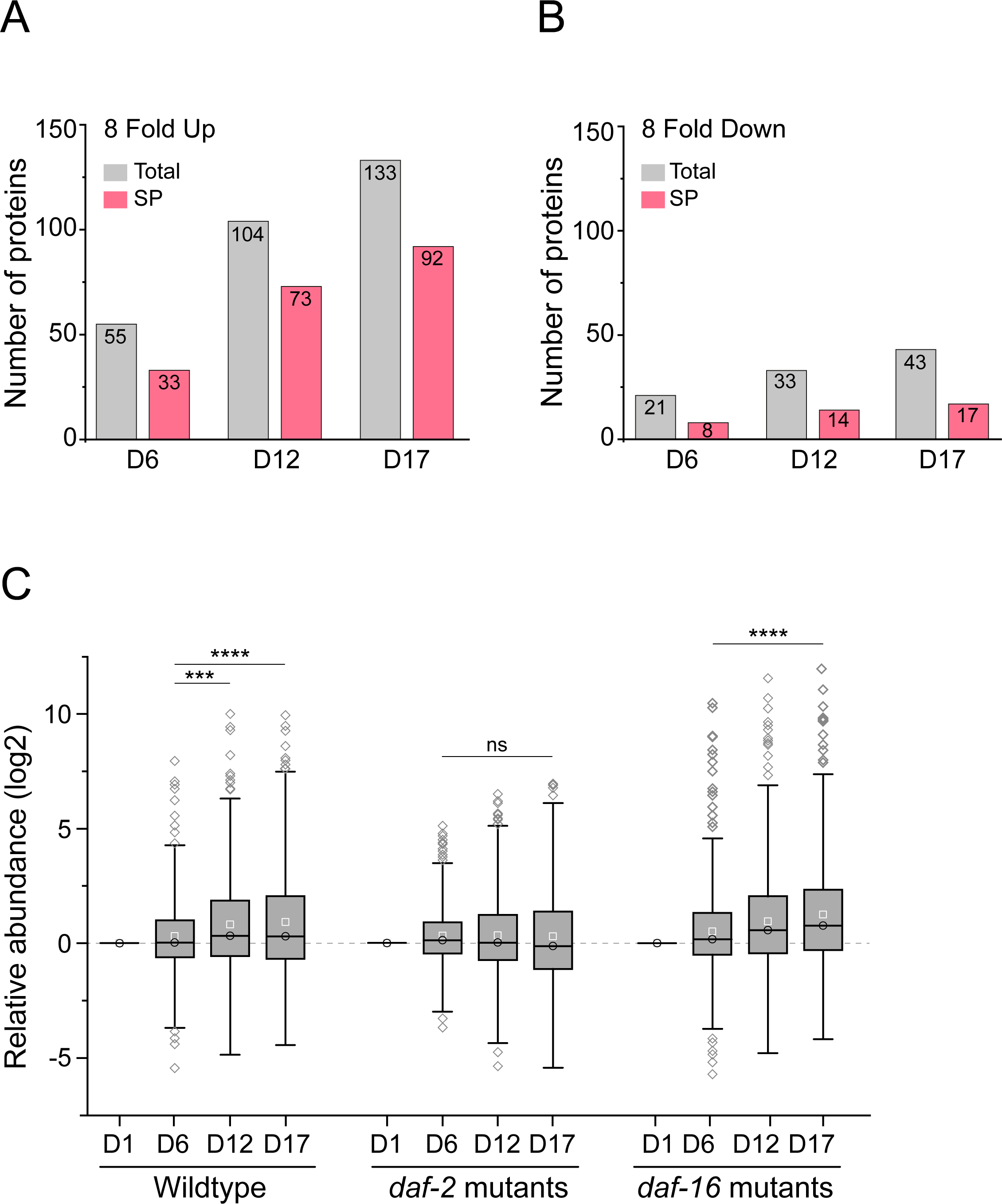
Abundance analysis of the SP proteins during aging. **A-B** Bar chart representing the number of upregulated and downregulated (greater than 8-fold) SP proteins that change with age in the Walther dataset. The grey color denotes total proteins while the pink color represents SP proteins. Number of proteins are plotted on the y-axis and time points in days mentioned on the x-axis. **C** Boxplots showing the relative abundance of the SP proteins (log2 values) in different lifespan mutants. In each lifespan mutant, the relative abundance level is calculated with respect to day1 values. The circles in the box represent the mean values whereas the squares are the median values. (P values for t-test: WT D6-D12: 0.004, WT D6-D17: <0.0001, *daf-2* D6-D12: 0.6949, D6-D17: 0.1387, *daf-16* D6-D12: <0.0001, daf-16 D6-D17: <0.0001). (P values in Mann Whitney U test are <0.0001 for all). (* <0.05, **<0.01, ***<0.001, ****<0.0001).

In *C. elegans* insulin like signalling pathway regulate proteostasis and the lifespan (Hipp et al., 2019). We wanted to find whether the SP protein abundances are regulated in the lifespan mutant worms. To do this we looked at abundance levels of the SP proteins in *daf-2* (long-lived) and *daf-16* (short-lived) mutant worms comparing with wildtype worms. Our analysis indicated that the abundance changes of the SP proteins during aging in the lifespan mutant worms follow same trend as wildtype worms. However, the mean abundance is slightly less for the *daf-2* and more for the *daf-16* mutant worms when compared to the young worms of the same strain or mean of the age matched wildtype worms (Fig. 3C). Overall, our results showed that the abundance of the SP proteins increase more than the decrease and lifespan mutant differently regulate these abundances during aging.

### 3.4. Aggregation of the SP proteins during aging

Protein aggregation is inherent property of aging and increases with age. We wanted to know whether the abundance increase of the SP proteins result in their aggregation with age. To found this, we analysed age related protein aggregation data from Walther et al (Walther et al., 2015). First, we analysed all the proteins identified in Walther et al aggregation data (WT, day12 vs day1) and selected the SP proteins (Fig. 4A). As shown in the volcano plot, 134 SP proteins identified in the aggregated proteins (red circles) and majority of them have higher aggregation abundances. This result indicates that with age, the SP proteins aggregate and their abundance in the aggregation increases. It has been reported that high abundance proteins also contribute more to aggregation load. To check abundance of these 134 SP proteins, we compared their total abundance with their aggregation abundance at day12 in wildtype worms. There was a positive correlation (Fig. 4B), suggesting that the aggregated SP proteins are also have higher total protein abundances. When protein abundances increase more than their critical solubility, they become supersaturated and result in their aggregation (Ciryam et al., 2015). Our results indicate that supersaturation of a subset of the SP proteins result in their aggregation during aging.

**Fig. 4.**
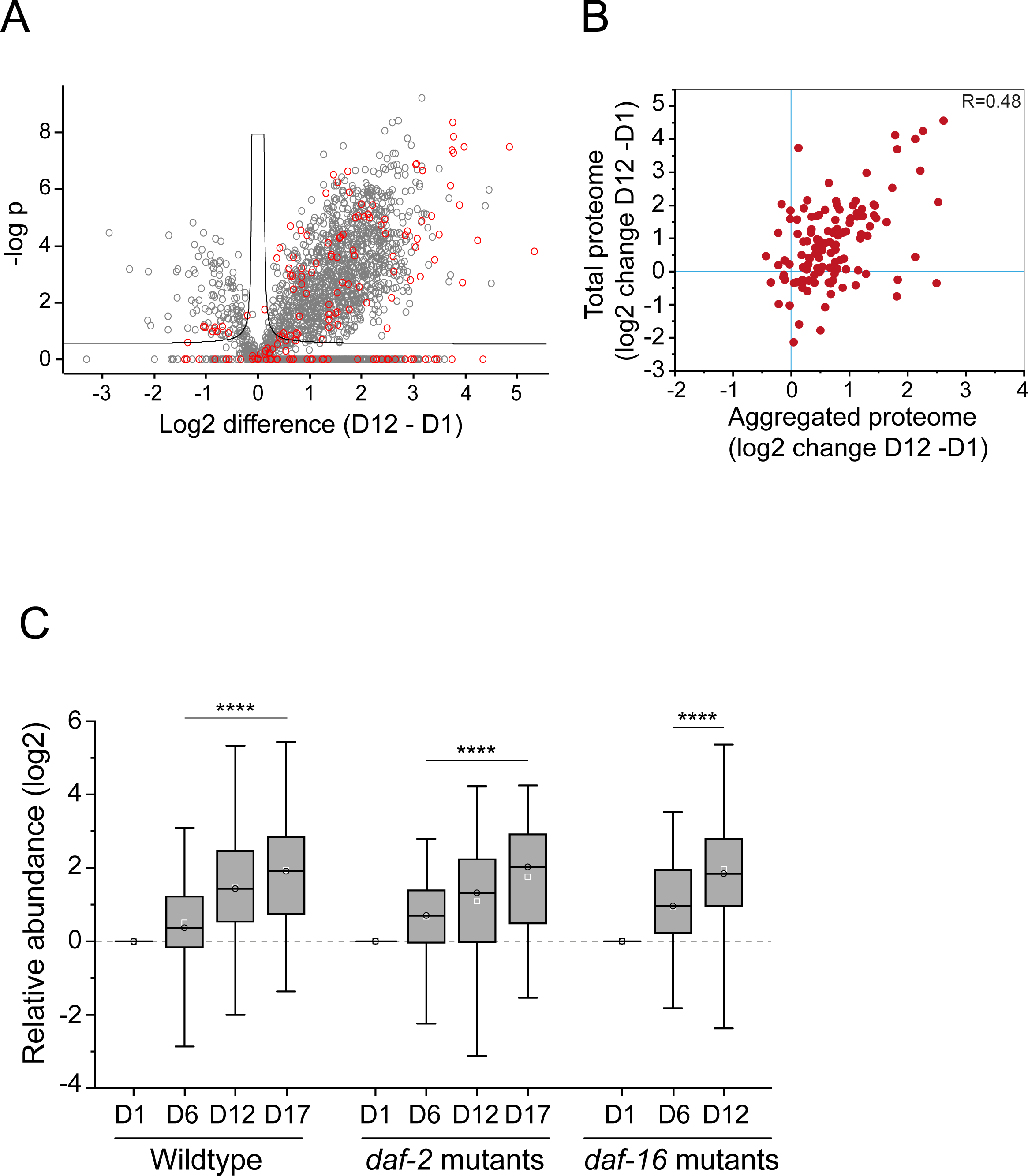
Aggregation of the SP proteins during aging. **A** Volcano plot showing aggregation of day12 WT worms (relative to day1). All the identified protein were filtered based on at least three valid values. The grey points correspond to total identified proteins (1966), the red points correspond to the SP proteins (134). The X-axis represents the log2 difference of day12 compared to day1 and Y-axis represents the −logP value. **B** Positive correlation between abundance changes in the aggregated proteins and abundance changes for the same proteins in the total proteome. Abundance differences is for day12 of WT worms. The Pearson correlation coefficient R is indicated. **C** Boxplots representing the relative abundance of total aggregation (log2 values) in different lifespan mutants. In each lifespan mutant, the relative abundance level is calculated with respect to day1 values (plotted on y-axis). The circles in the box represent the mean values whereas the squares are the median values. Time points in days and strains are mention on the x-axis. (P values: WT D6-D12: <0.0001, WT D6-D17: <0.0001, *daf-2* D6-D12: <0.0001, D6-D17: <0.0001, *daf-16* D6-D12: <0.0001). (* <0.05, **<0.01, ***<0.001, ****<0.0001).

Next, we asked whether aggregation of the SP proteins is modulated in the lifespan mutant worms. We looked at aggregation of the SP proteins in *daf-2* and *daf-16* mutant worms and compared with the WT. We found that the aggregation of SP proteins increases with age in all the three strains (Fig. 4C and Table S4). However, there is a modest difference in the mean abundance at day12, where it is slightly less in *daf-2* and more in *daf-16* mutant worms compared to WT. This result can be correlated with the total abundance levels of the SP proteins in the lifespan mutants. This analysis suggests that abundance level and aggregation of a subset of the SP proteins are modulated in the *daf-2* mutant worms in a way balance the proteostasis.

### 3.5. Overlap of the SP proteins with extracellular vesicles (EVs) proteins

Extracellular vesicles (EVs) are membrane bound particles derived from various cells. They move around in the body fluids or extracellular fluids carrying bioactive molecules such as proteins, lipids, and nucleic acids to mediate intercellular communication (van Niel et al., 2018). It has been observed that senescent cells release more and altered EVs compared to nonsenescent cells, and these senescent EVs can induce senescence in other healthy cells (Yin et al., 2021). There is also growing evidence that EVs may be directly involved in the aging process by mediating cell nonautonomous signaling. The age induced alteration in EVs thereby contributing to a degradation in intercellular communication is considered a hallmark of aging (Manni et al., 2023). To check whether the SP proteins in this analysis are part of EVs, we made a comparison with publicly available *C. elegans* EV proteome data by Russell et al and Nikonorova et al (Russell et al., 2020; Nikonorova et al., 2022). Together, these two studies identified EV cargo (proteins and RNA) shed by *C. elegans* (young males and hermaphrodites) into the environment.

Out of the 916 SP proteins, we found presence of 206 SP proteins in the EVs from Russell data (Fig. 5A) and 462 SP proteins in the Nikonorova data (Fig. 5C, Table S5). This suggest that subset of the age regulated SP proteins move to extracellular space in younger age to perform their putative functions in the extracellular space or as signalling molecules. Since abundance of the many SP proteins increase with age, we wanted to check whether the SP proteins also follow the trend. We analysed abundance levels for these EV associated proteins in the WT as well as in the lifespan mutant worms. We found that majority of the EV associated SP proteins also continue to increase their abundance with age (Fig. 5B and D). Intriguingly, this abundance is less pronounced in *daf-2* mutant worms compared to WT. In contrast, this abundance increase is more pronounced in the *daf-16* mutant worms. These results suggest that the abundances of a subset of EV proteins is differentially regulated in the lifespan mutant worms. We assume that the *daf-2* mutants worms maintain lower abundance levels of a subset of the SP proteins and/or EV associated proteins to maintain their function and balance the proteostasis in extracellular space.

**Fig. 5.**
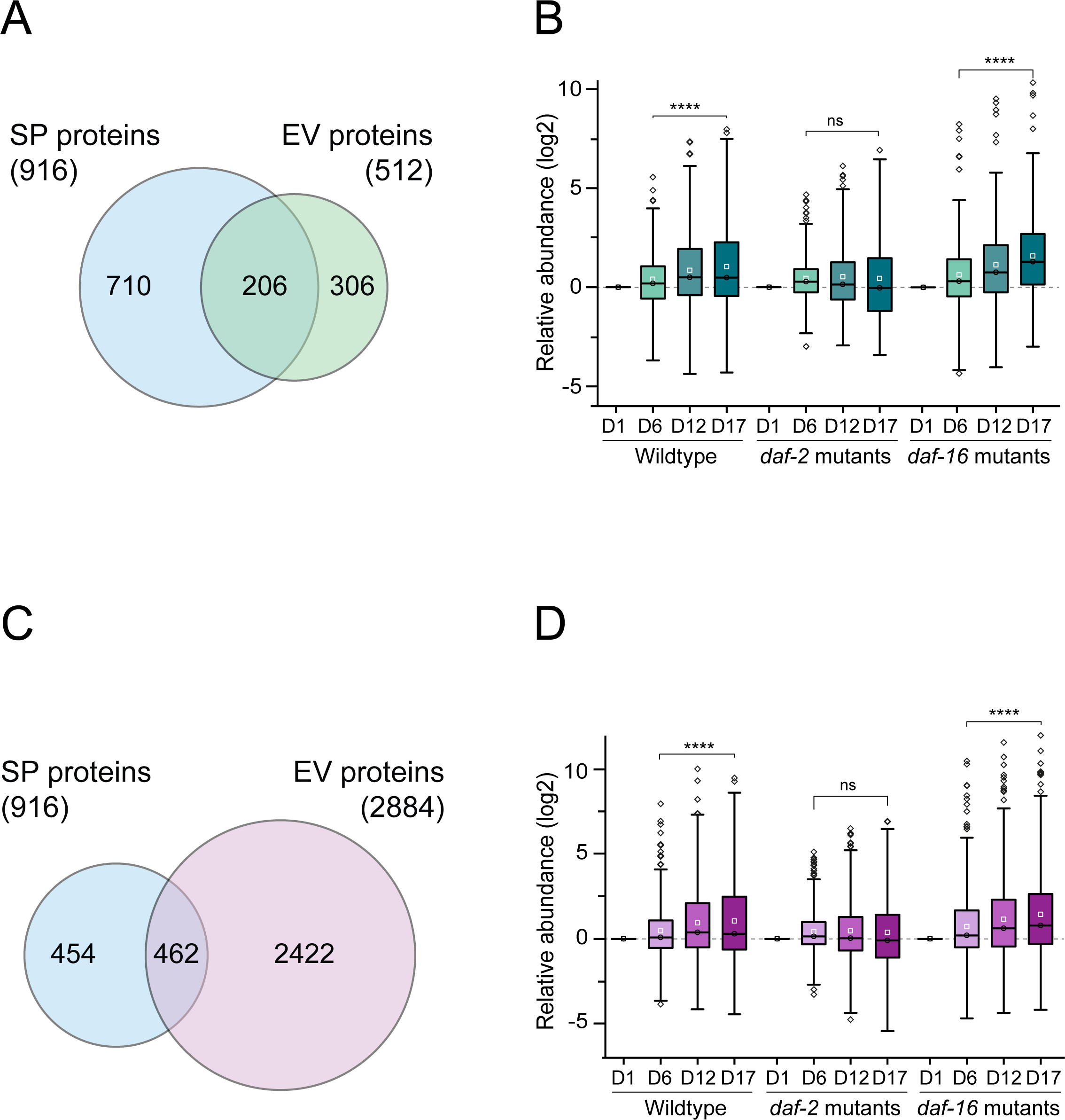
Overlap of SP proteins with extracellular vesicles (EVs) proteins. **A** Venn diagram illustrating the number of identified EV Proteins and the common ones with Russell et al data. **B** Boxplots showing the relative abundance of the SP proteins associated with the EVs in Russell et al data in different lifespan mutants. The relative abundance level is calculated with respect to day1 values (plotted on the y-axis). Time points in days and strains are mention on the x-axis. **C** Venn diagram illustrating the number of identified EV Proteins and the common ones with Nikonorova et al data. **D** Boxplots showing the relative abundance of the SP proteins associated with the EVs in Nikonorova et al data in different lifespan mutants. The relative abundance level is calculated with respect to day1 values (plotted on the y-axis). Time points in days and strains are mention on the x-axis.

### 3.6. Abundance of the secreted proteins limits the lifespan

Since we found differential abundances for a subset of the SP proteome, we selected four proteins (two with human orthologs and two nematode specific) based on their abundance levels in the lifespan mutants and wanted to check their effect on lifespan of WT worms. To check effect of the higher abundance and aggregation of the SP proteins on lifespan, we selected two proteins with human orthologs (W01F3.2 and K08D8.6) and two nematode specific proteins (Y62H9A.5 and Y62H9A.6) that differentially expressed in lifespan mutant worms compared to WT (Table S6A). Y62H9A.6 or *ule-5* has already shown to be secreted and abundance increase in old age (Zimmerman et al., 2015). First, we checked their total abundance and aggregation during aging. We found that abundance of these proteins is less in the *daf-2* and more in *daf-16* mutant worms compared to WT (Fig. 6A). This suggest that lower levels of these proteins might beneficial to WT worms. To test whether the high abundance levels of the SP proteins limit the lifespan in WT worms, we knocked-down the genes coding for these proteins by RNAi. Compare to the empty vector control, knocking down of W01F3.2 resulted in 25%, K08D8.6 in 20%, Y62H9A.5 in 19% and Y62H9A.6 in 16% increase in lifespan of WT worms (Fig. 6B-D and Table S6B). Knocking down *daf-16* decreased lifespan by on average 20%, validating our RNAi experiments. Our results showed that reducing levels of the high abundant proteins had a positive effect on lifespan, suggesting that high abundance of SP proteins might limit the lifespan.

**Fig. 6.**
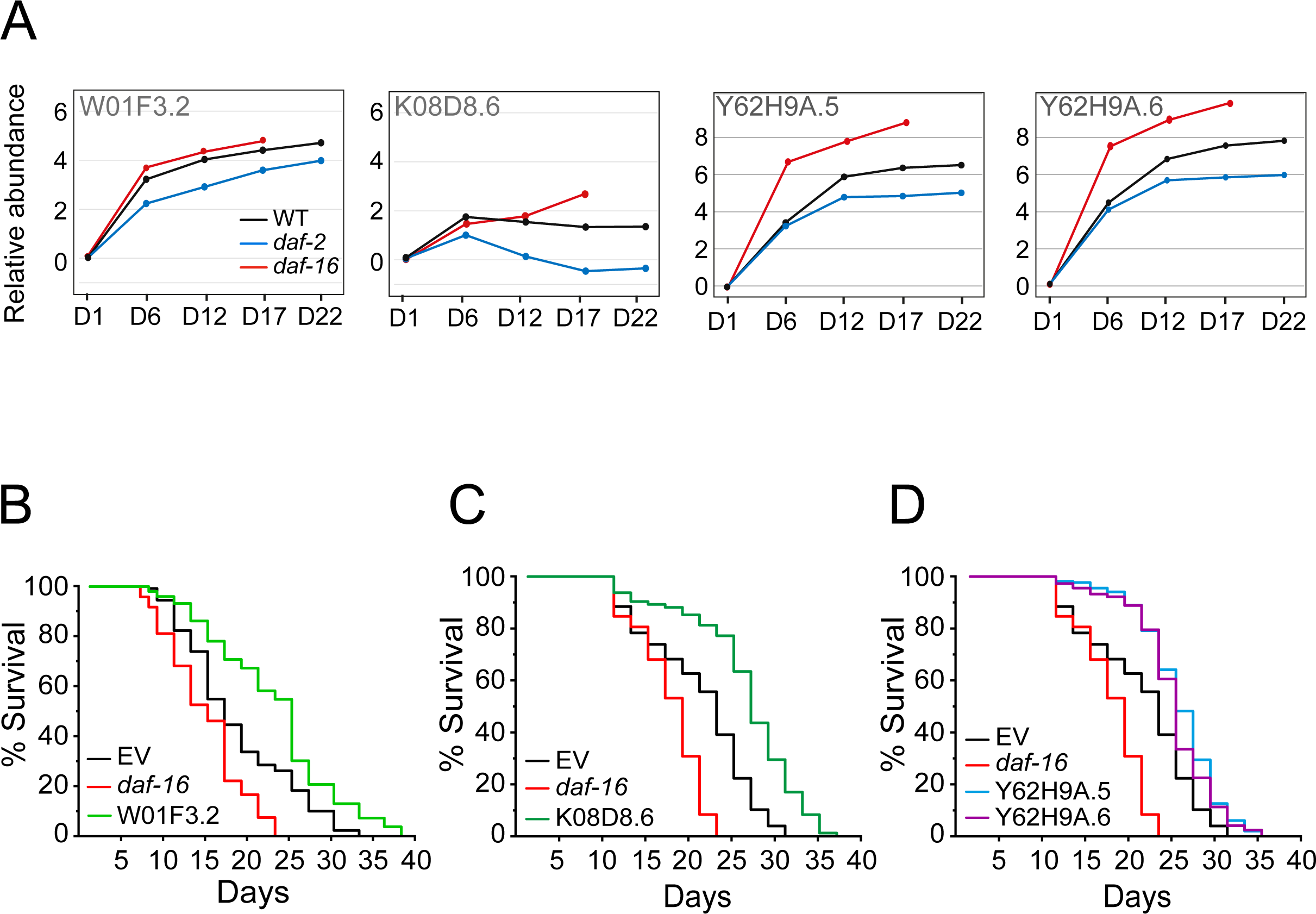
Lifespan Analysis of the selected candidates. **A** Protein levels of identified secreted proteins with age in different WT and lifespan mutant worms. The log2 fold changes are related to day1 (plotted on the y-axis). Time points in days are mention on the x-axis. The black line denotes WT (N2), the blue line represents *daf-2* (long-lived mutant) and the red line represents *daf-16* (short-lived mutant). **B-D** Lifespan curves showing that RNAi of W01F3.2, K08D8.6, Y62H9A.5 and, Y62H9A.6 show an increment in lifespan in comparison to WT and short-lived mutants.

## 4. Discussion

In multicellular organisms, intercellular communication is essential for many cellular functions to maintain organism wide homeostasis. Secreted proteins play key roles in these cell-nonautonomous processes and are found to be increased in response to stress, during infections and aging (Liang et al., 2014; Walther et al., 2015; Narayan et al., 2016; Suh and Hutter, 2012). These secreted proteins need to maintain their functional states once they released to the extracellular space. Otherwise, the organism homeostasis gets imbalanced and can lead to pathological conditions (Satapathy and Wilson, 2023; Yuyama and Igarashi, 2016). Identifying and characterizing stress/age induced extracellular proteins provide onset of diseases and opportunities for biomarker discovery.

In *C. elegans* ∼17% genes are predicted to encode secreted proteins and many of these are uncharacterized (Suh and Hutter, 2012). Here, we present an analysis of age-related changes in SP proteins that exhibit potential to be localized to extracellular space. We identified 916 SP proteins from two studies (Walther et al., 2015; Narayan et al., 2016), in which a subset of the proteins is associate with extracellular vesicles. We show that abundance increase is more prominent than the decrease during aging. The PPI and network analysis revealed that the age-regulated SP proteins involve in multiple biological process such as metabolic, lipid binding, ion transport and protein folding. However, fatty acid binding and endopeptidase inhibitors are associated with upregulated proteins whereas endopeptidase activity and unfolded protein binding are associated with downregulated proteins. This implies that with age protein degradation impairs in the extracellular space and unfolded proteins increase, which result in abundance increase of the SP proteins. This abundance increases once reach its critical solubility result in aggregation of the proteins. Indeed, many of the SP proteins aggregate with age and their abundance in the total and aggregation also increase. A few of the analysed SP proteins from this study were already shown to be secreted with increased abundance during aging and aggregation supporting our prediction (Zimmerman et al., 2015; Gallotta et al., 2020).

Since some SP proteins can secrete with extracellular vesicles (EV) and content of the EVs from senescent cells is different from healthy cells, we looked for the SP proteins in the EVs secreted from young *C. elegans*. Abundance and aggregation of these EV associated proteins also increase with age. EV release is crucial for development, but overproduction can disrupt it, as shown in *C. elegans* studies. Therefore, EV release must be tightly regulated to ensure the correct number of EVs are produced at the right time (Beer and Wehman, 2017). As we found abundance increase for some of the EV associated proteins with age, we assume uncontrolled production of EVs during aging may alter their secretion pattern. This may result accumulation of these proteins inside the cells if secretion is hindered. Alternatively, the EVs can still secreted out and they may get accumulated in the extracellular space. The follow up studies could be designed to test this. We speculate that some of the SP proteins may function only during development and they may not require post reproduction and their accumulation result in their loss of functions that limit the lifespan.

To better understand fates of these SP proteins and EV associated SP proteins with age, we monitored them during aging in the lifespan mutant worms. As expected, their abundance level is maintained lower in the long-lived mutant worms compared WT. However, abundance levels for the same proteins are more in *daf-16* short-lived mutant worms compared to WT, suggesting that the higher abundance levels reduce lifespan. In agreement with, reducing levels of some secreted proteins increase lifespan and over expression shorten lifespan (Zimmerman et al., 2015; Ezcurra et al., 2018; Seah et al., 2016). To further validate we selected two proteins with human orthologs and two nematode specific proteins. Our findings also confirm that elevated levels of extracellular proteins restrict lifespan, hinting that long-lived mutant worms regulate them differently to maintain better proteostasis. As certain identified extracellular proteins in this analysis share orthologs with humans, there is an intriguing opportunity to investigate selected proteins in samples from human aging or disease contexts.

## Acknowledgements

The authors acknowledge BioX centre, SBB at IIT Mandi for providing facilities.

## CRediT authorship contribution statement

Prasun Kumar: Data curation, Formal analysis, Investigation, Methodology, Validation, Visualization, Writing – review & editing. Vishwajeet: Data curation, Formal analysis, Investigation, Methodology, Validation, Visualization. Prasad Kasturi: Conceptualization, Data curation, Formal analysis, Funding acquisition, Methodology, Supervision, Visualization, Writing – original draft, Writing – review & editing.

## Funding

This work was supported by funding from Department of Biotechnology, Government of India, as DBT-Ramalingaswami Re-entry grant (BT/RLF/Re-entry/31/2018 to P.K).

## Declaration of Competing Interest

The authors have no conflicts of interest to report.

## Appendix A. Supporting information

The datasets used in this work are available in the supplemental file.

---

**Supplementary Fig. 1:**
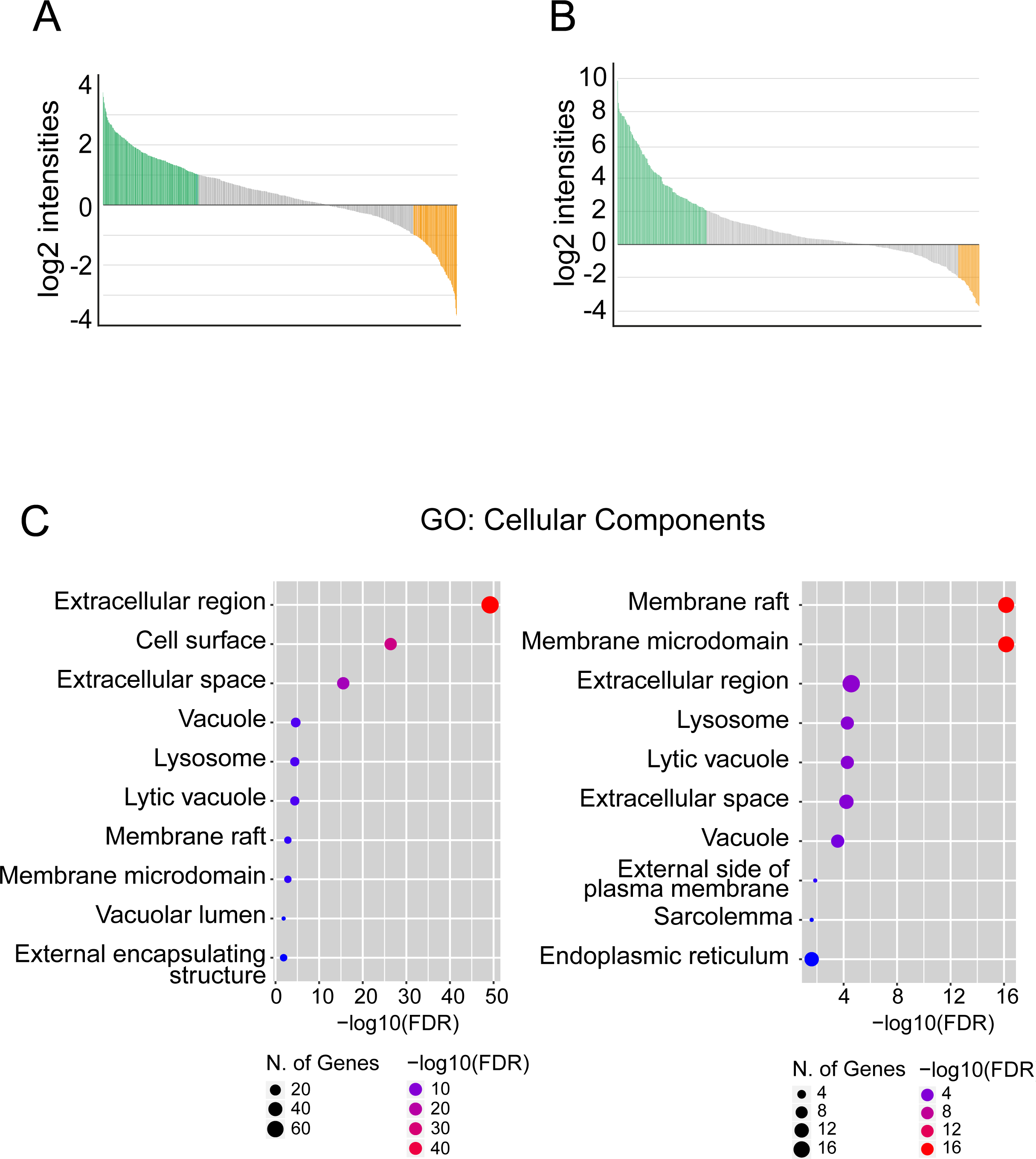
Age related changes in SP proteins. **A** Overview of the dynamic range of the SP proteins at day10 (Narayan data) quantified using log2 intensities. The green color represents upregulated proteins, and the orange color denotes downregulated more than log2 intensities of 2 respectively. **B** Overview of the dynamic range of the SP proteins at day10 (Narayan data) quantified using log2 intensities. The green color represents upregulated proteins, and the orange color denotes downregulated more than log2 intensities of 2 respectively. **C** Enriched gene ontology (GO) cluster of cellular components (CC) of the up and down regulated SP proteins identified both in the Walther and Narayan datasets. Left, upregulated (log2 > 1.5) (401 entries) and right, downregulated (log2 < 0.7) proteins (202 entries).

**Supplementary Fig. 2:**
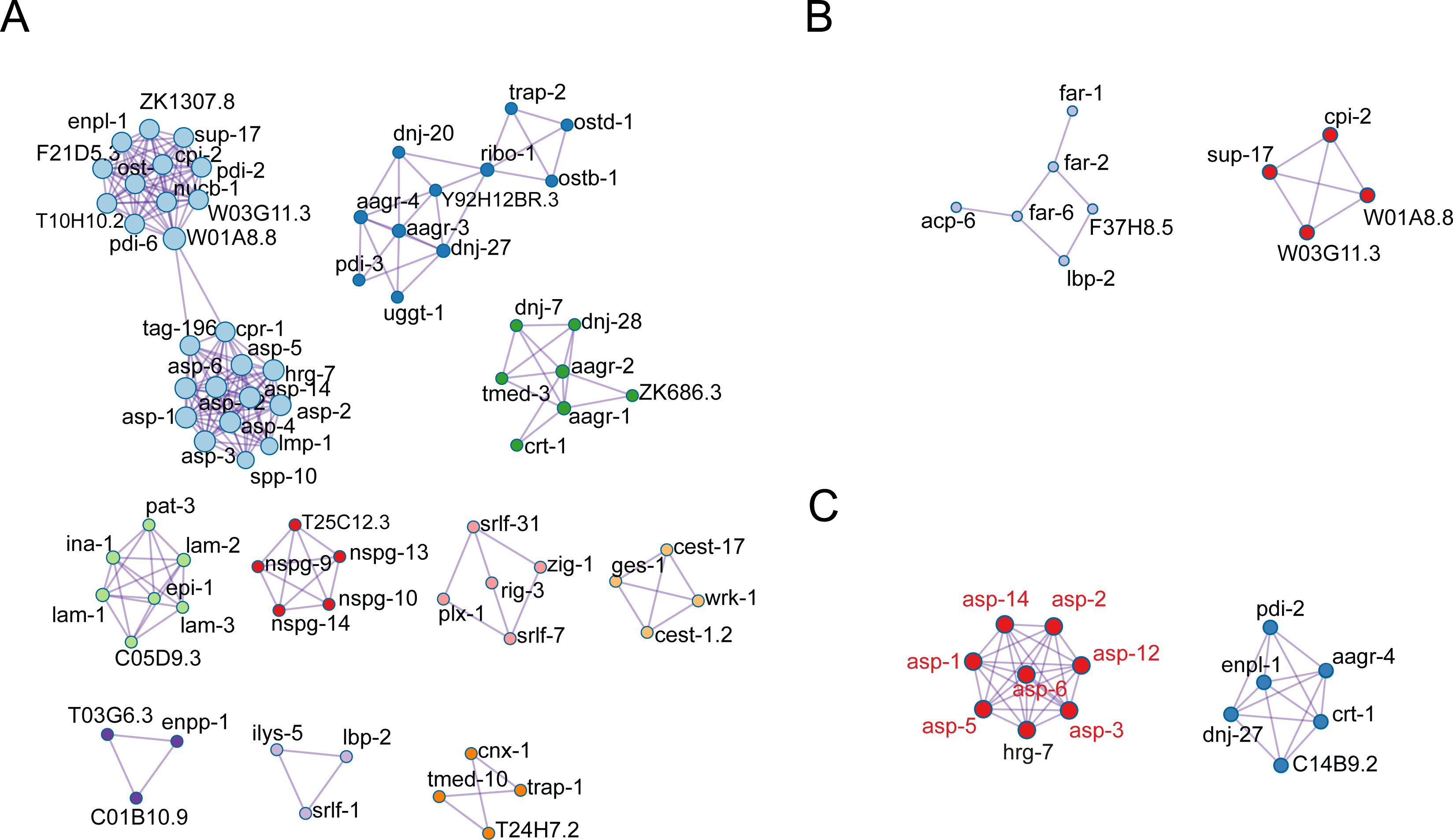
PPI network of SP proteins. **A** Protein-Protein Interaction (PPI) Network showing more prominent interactions (based on log10 P values) between all identified SP proteins. **B** Protein-Protein Interaction (PPI) showing more prominent interactions (based on log10 P values) for the SP proteins upregulated by 1.5-fold. **C** Protein-Protein Interaction (PPI) showing more prominent interactions (based on log10 P values) for the SP proteins downregulated by 1.5-fold.

**Supplementary Fig. 3:**
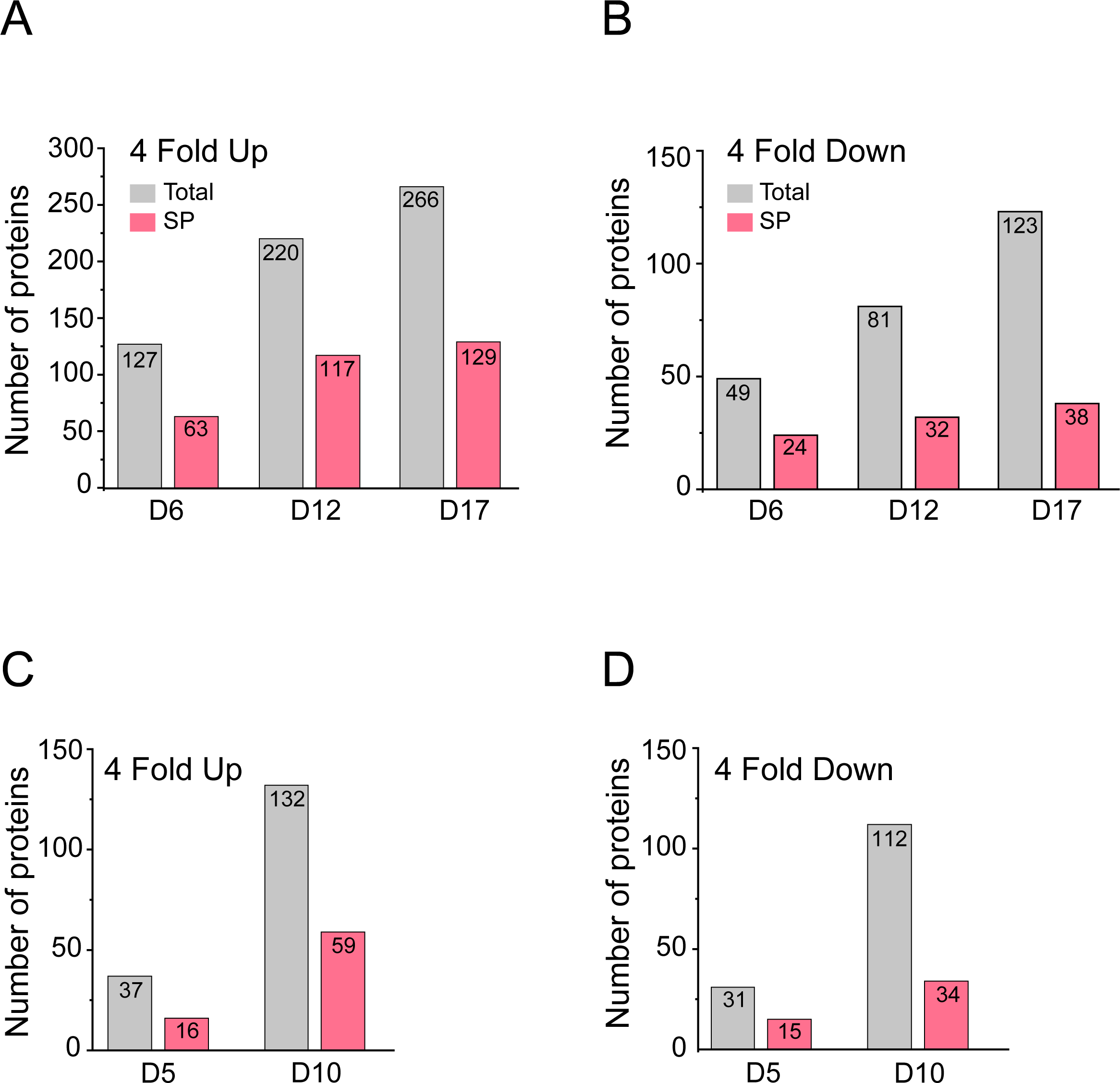
Abundance levels of SP proteins. **A-B** Bar chart representing the number of Upregulated and Downregulated (Greater than 4-fold) SP proteins that change with age in the Walther dataset. The grey color denotes total proteins while the pink color represents SP proteins respectively. **C-D** Bar chart representing the number of Upregulated and Downregulated (Greater than 4-fold) SP proteins that change with age in the Narayan dataset. The grey color denotes total proteins while the pink color represents SP proteins respectively. Number of proteins are plotted on the y-axis and time points in days mentioned on the x-axis.

## Notes

### Competing Interest Statement

The authors have declared no competing interest.

